# SNP calling for the Illumina Infinium Omni5-4 SNP BeadChip kit using the butterfly method

**DOI:** 10.1101/2022.01.17.476594

**Authors:** Mikkel Meyer Andersen, Steffan Noe Christiansen, Jeppe Dyrberg Andersen, Poul Svante Eriksen, Niels Morling

## Abstract

We introduce the “butterfly method” for SNP calling with the Illumina Infinium Omni5-4 BeadChip kit without the use of Illumina GenomeStudio software. The method is a within-sample method and does not use other samples nor population frequencies to call SNPs. The butterfly method is based on a three-component mixture of normal distributions, in which parameters are easily found using the open-source statistical software R. This makes the method transparent, straight-forward to change parameters according to the user’s needs, and easy to analyse the data within R after the SNPs have been called. We contribute with two open-source R packages that make SNP calling easy by helping with bookkeeping and by giving easy access to meta-information about the SNPs on the Illumina Infinium Omni5-4 BeadChip Kit (including chromosome, probe type, and SNP bases). We test our method on > 4 mio. SNPs and compare the results with those obtained with the GenTrain method used by Illumina GenomeStudio as well as SNPs obtained by PCR-free whole genome sequencing (WGS). We demonstrate two variants of our method: one where we account for potential probe type bias by estimating a separate model for each probe type (type I and type II) and another that uses a general model such that the model’s parameter estimates do not depend on the sample that is being analysed. We focused on varying the no-call rate and show how it changed the concordance with that of WGS. This is done by using a threshold on the *a posteriori* probability of belonging to a SNP cluster and by using the number of beads to adjust the stringency of the no-call mechanism. With the butterfly method, we achieve a SNP call rate of around 99% and a SNP concordance of around 99% with the WGS data. By lowering the *a posteriori* probability threshold for no-calls, we can get a higher call rate fraction than the GenomeStudio and by using a higher *a posteriori* probability threshold, we can achieve a higher concordance with the WGS data than the GenomeStudio.

## 1. Introduction

We introduce a method for SNP calling that is exemplified with the Illumina Infinium Omni5-4 SNP BeadChip kit [1], but it can potentially be used with other Illumina BeadChip SNP kits. The Omni5-4 is based on the Illumina BeadChip technology that has two types of probes, type I and type II. The nucleotides are detected by two colour channels: red (detects A/T nucleotides) and green (detects C/G nucleotides). Probe type I has two bead addresses such that one nucleotide is interrogated by the beads with one address and the other nucleotide by the beads with the other address. The light emitted by the fluorophores attached to the bases are measured by the same colour channel on both bead addresses (by design of probe sequence), i.e., either red or green. The other colour channel is not used for this probe type (but can potentially be used for estimating e.g., background noise). Probe type II has only one bead address, such that the signal from one nucleotide is measured by the red colour channel and the other nucleotide is measured by the green colour channel. The output from the array is the mean signal intensity from each colour channel together with the standard deviation and the number of beads investigated.

This technology can give two alleles that – to some confusion – are called A/B alleles [2, 3]. Simplified, the A/B system tells whether the SNP position has homozygous reference alleles, homozygous alternative alleles, or heterozygous alleles. The A/B alleles must be converted to the standard DNA bases A, T, G, and C using a manifest file [4] from Illumina.

Bookkeeping with the selection of the right colour channel for the right probe and conversion of the A/B allele system to the standard DNA bases is a tedious task with many different rules that must be applied. We have developed an open-source software that can help to do these tasks. The software is published as an R [5] package called `snpbeadchip` [6] for selecting the correct colour channels and converting the A/B alleles to plus/minus alleles and an accompanying R package called omni54manifest [7] that provides easy access to the information about the probes such as the manifest [4] and mapping information [8]). `snpbeadchip` uses `illuminaio` [9] to read idat files.

Once the signal intensities of the reference and alternative allele are obtained, the SNP can be called. We propose a method that we refer to as the “butterfly method” (cf. Fig. 3).

We tested the butterfly method against high quality PCR-free WGS data, which are considered the gold-standard for “concordance”. Furthermore, we compared the butterfly method with SNP calls from the Illumina GenomeStudio software [10].

The aims of this paper and the purpose of the method is to provide an open description of a SNP calling method that others can use, replicate, and improve. The description of the method used by GenomeStudio called GenTrain 3.0 [11] is not publicly available [12]. It uses the data of the other samples for the sample analysis, which can be problematic because the SNP calling of one sample is influenced by the other samples in the analysis. One such problematic situation may arise if the samples analysed together are of varying quality, e.g., the combination of samples with high quality and partly degraded DNA, as may be the case in a forensic setting.

## 2. Materials and Method

All analysis were made using R [5] version 4.1.2 and tidyverse [13].

### 2.1. Blood samples and DNA extraction

Peripheral blood from four individuals was collected and stored at −20^◦^C until DNA extraction. DNA extraction was carried out using the DNeasy Blood & Tissue Kit (Qiagen) following the manufacturer’s recommendations for purification of total DNA from whole blood.

### 2.2. SNP typing using Illumina Infinium Omni5-4

All samples were analysed using the Illumina Infinium Omni5-4 kit following the manufacturer’s recommendations with varying DNA input amounts. The DNA concentration was measured using the Qubit dsDNA HS Assay Kit (Thermo Fisher Scientific). Two-fold serial dilutions of DNA from three samples were performed using nuclease-free water to obtain samples with the following DNA amounts: 400 ng, 200 ng, 100 ng, 50 ng, and 25 ng. The DNA amount of the fourth sample was 400 ng. Briefly, the DNA was hybridised to the probes attached to the BeadChips. Hereafter, the attached probes were subject to singlebase extension and stained. The BeadChips were scanned using the iScan^™^ system (Illumina) following the manufacturer’s recommendations

### 2.3. PCR-free whole genome sequencing

PCR-free WGS and variant detection were carried out as described in [14].

### 2.4. The butterfly method

“The butterfly method” i s based on a finite mixture of bivariate normal distributions [15] with three mixture components (one for each SNP/genotype, i.e., AA, AB, or BB in the A/B allele system [2]).

Let *A* and *B* be the mean signal intensities for allele A and B in the A/B allele system, respectively. We log transformed (using the natural logarithm) the mean signal intensities with one added (to avoid numerical problems as the mean signal intensity can be 0, and ln(0) = −∞). Thus, we used *A*′ = ln(*A* + 1) instead of *A* in the models.

Using *A*′ = ln(*A* + 1) and *B*′ = ln(*B* + 1), the model specifies a joint probability density function by

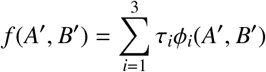

where *i* ∈ {1, 2, 3} indicates SNP group (e.g., *i* = 1 means AA, *i* = 2 means AB, and *i* = 3 means BB), φ_*i*_ is a probability density function for a bivariate normal distribution, and τ*_i_* = *P*(*i*) is the *a priori* (without taking intensities *A*′ and *B*′ into account) probability that the SNP has type *i* (and 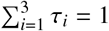).

In other words, we model the signal intensities as a three-component mixture of bivariate normal distributions. The signal intensities *A*′ and *B*′ can either come from SNP group 1 (AA), 2 (AB) or 3 (BB). The likelihood of observing *A*′ and *B*′ in each SNP group is weighted by τ_*i*_.

The unknown parameters in the model include e.g., the *a priori* probabilities, τ_*i*_, the mean values, and covariance matrices for the bivariate normal distributions (not shown). The parameters were estimated using the R package mclust [15]. We chose to model the mixture components as bivariate normal with any shape and orientation; in the mclust terminology this is called a `VVV` model.

When calling SNPs, we want to calculate the *a posteriori* probability of SNPs belonging to SNP group *k* given the signal intensities *A*′ and *B*′, which is given by

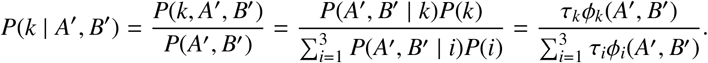

This is the probability of being in group *k* regardless of the allele intensities *A*′ and *B*′, τ_*k*_, multiplied by the likelihood of the allele intensities in SNP group *k*, *ϕ*_*k*_(*A*′, *B*′), and normalised (denominator) so that the *a posteriori* probabilities over the three SNP groups sum to 1.

We present three variants of the butterfly method. Different data sets were used to train (estimate the parameters of) the three-component mixture model: 1) each sample was its own reference using all SNPs simultaneously, 2) like 1, but using separate models for the two probe types (I/II), and 3) an ensemble model using all samples to estimate a single model.

#### 2.4.1. Calling SNPs

For the WGS data, we only used biallelic SNPs with a read depth of at least 25. We used the recommended settings in GenomeStudio [10].

For the butterfly method, we called the SNP with the maximal *a posteriori* threshold, except for situations with no-calls (NC). We chose to always make a NC if the mean signal intensities for both allele A and B were 0.

We investigated two ways of making a NC. Firstly, if the maximal *a posteriori* probability was below a certain threshold, we made a NC. This was done for a range of thresholds (from 0.5 to 0.999). Secondly, we chose to consider the number of beads with which the SNPs had been investigated, and the mean signal intensities were based on. If the number of beads was below five, we made a NC. We also used a threshold of zero beads.

For probe type II, the same beads capture both alleles, so there is only one number of beads for each investigated position. For probe type I, different beads capture each allele, so there is a number of beads for allele A and another for allele B. For probe type I, both numbers of beads must be above the threshold.

Imposing such NC thresholds results in calling fewer alleles but with higher confidence in the alleles called.

#### 2.4.2. Other methods

The GenoSNP method introduced in [16] is a within-sample method that uses a four-component mixture of *t*-distributions (like the normal distribution, but with heavier tails), where the forth cluster is a “null class” for capturing outliers. The calls are made by identifying the cluster with the maximal *a posteriori* probability, and hence, the no-calls are selected when the null class has the highest *a posteriori* probability. This is problematic as all outliers do not behave in the same way. Outliers are not expected to be distributed according to a *t*-distribution and grouped in the same cluster in the (*A*′, *B*′) space.

The M3 method introduced in [17] is similar to that in [16], except that a four-component mixture of normal distributions is used and the focus is on the ability to call rare variants. In [17], the *a posteriori* probability is mentioned, but only in connection with calculating the average *a posteriori* probability for each SNP.

To summarise, our paper contributes with the following novel work: a) analysis of data obtained with the Illumina Infinium Omni5-4 Kit by comparing SNP calls made by GenomeStudy and GenTrain 3.0 [10, 11]; b) demonstrating how *a posteriori* probabilities and the numbers of beads can be used to categorise NC (instead of including a less flexible null-cluster) and analyse how they impact the concordance with WGS calls; c) showing how a non-sample specific, a general model, and a sample and probe type specific model performs. Our method is available as the R software packages `snpbeadchip` [6], `omni54manifest` [7], and the existing R software package `mclust` [15] for estimating the mixture model (with the function `mclust`(…, G = 3, `modelNames` = “`VVV`”)) and calculating *a posteriori* probabilities (with the function `predict`()).

We chose not to include any of the above methods because the main focus of this paper was to explore the possibility of adjusting the NC rate and to offer open-source software for this purpose because none of the methods are supported by publicly available software.

## 3. Results

### 3.1. Included SNPs

The Omni5-4 manifest [7] has 4,327,108 SNPs. We removed 271,680 SNPs (details in Table 1) and ended up with 4,055,428 autosomal SNPs (93.7% of the original). Of the 4,055,428 SNPs included, 135,419 SNPs were typed with type I probes (ambiguous) and 3,920,009 were typed with type II probes (unambiguous).

**Table 1:** Filtering of probes. The filtering was performed sequentially in the order shown, and the numbers are conditional on the filtering order. Steps 2 and 4 are performed to remove probes with registered problems of different kinds.

| Step | Description | SNPs removed | SNPs left |
| --- | --- | --- | --- |
| 0 | Manifest file |  | 4,327,108 |
| 1 | Type II and ambiguous (rs28362918, rs28897688) | 2 | 4,327,106 |
| 2 | Chromosome 0 (probe problems) | 9,724 | 4,317,382 |
| 3 | Chromosome X/Y/mitochondria | 121,387 | 4,195,995 |
| 4 | Non-empty mapping comment (probe problems) | 636 | 4,195,359 |
| 5 | INDELs (insertion-deletions) | 4,412 | 4,190,947 |
| 6 | Multiple probes binding to same rsID | 129,739 | 4,061,208 |
| 7 | Multiple rsIDs for a name | 5,780 | 4,055,428 (93.7%) |

### 3.2. Number of beads

The density of the numbers of beads for both probe types for all four individuals are given in Fig. 1. For probe type I, the associations of the numbers of beads for alleles A and B are shown in Fig. 2.

**Figure 1:**
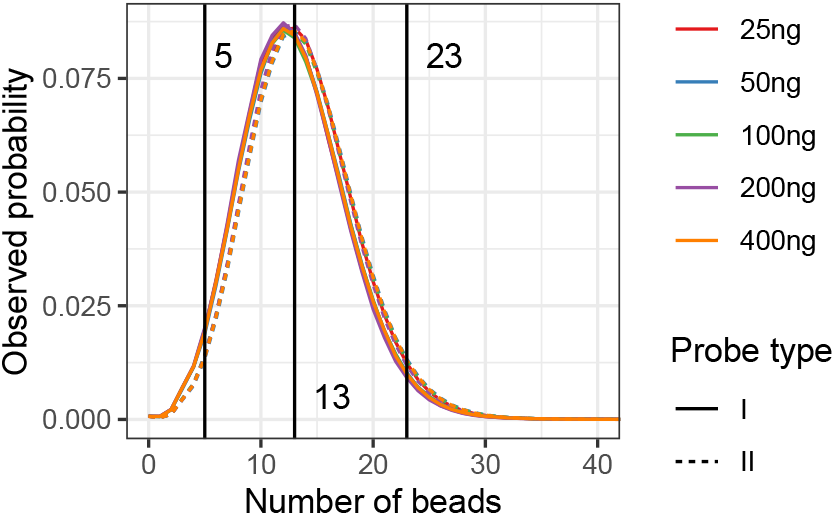
The density of the number of beads for both probe types for all four individuals. The interval [5, 23] is the middle 95% of the probability mass (this is the same for both probe types and all dilutions). Hence, for 95% of the probes it is expected that they have between 5 and 23 beads. The median number of probes for both probe types and all DNA concentrations was 13.

**Figure 2:**
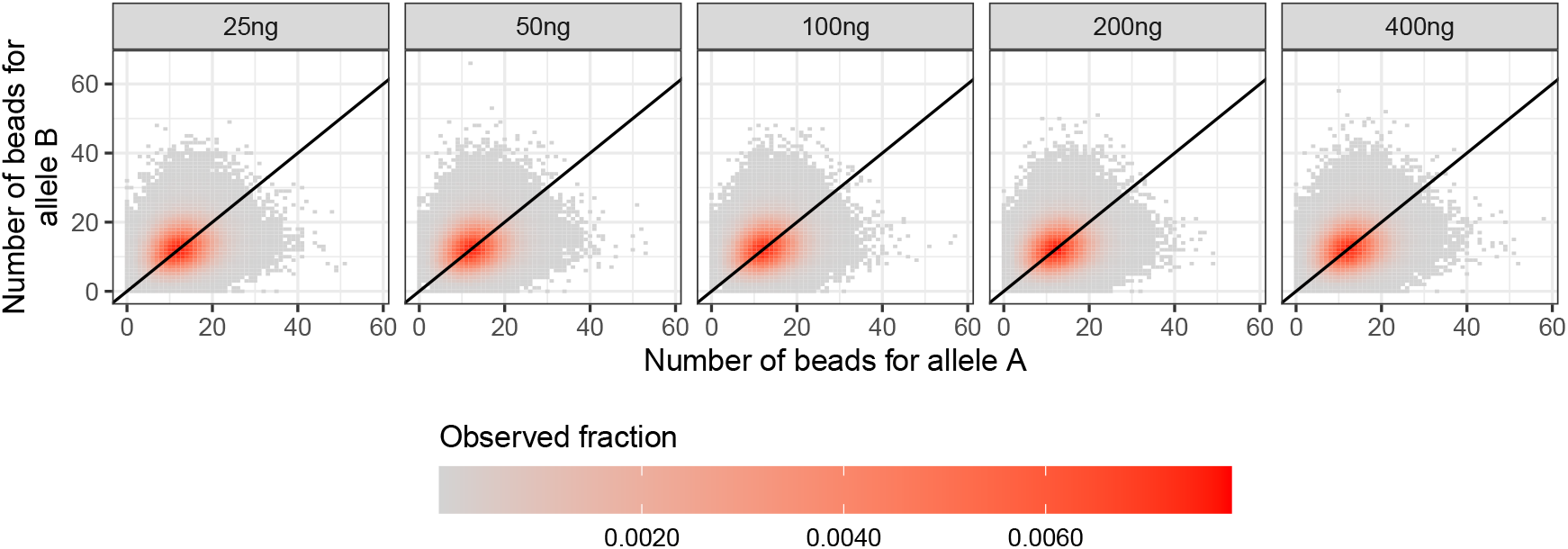
The association between the numbers of beads for allele A and B for probe type I.

### 3.3. Intensities

The mean signal intensities with an illustration of the models are shown in Fig. 3.

**Figure 3:**
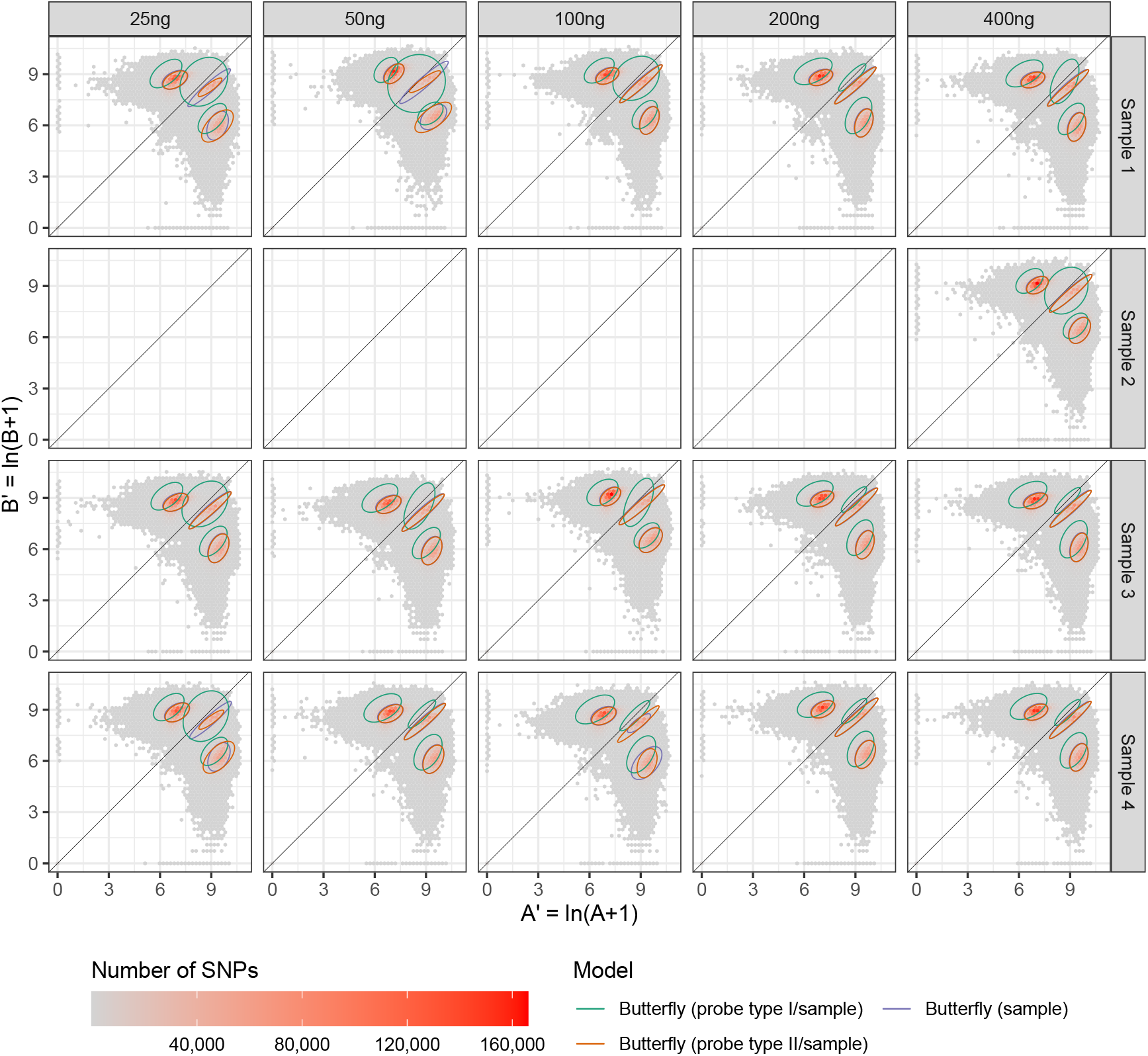
The mean signal intensities for allele A and B. The three SNP groups AA, AB, and BB (in A/B nomenclature) form butterfly patterns. The 75% confidence ellipses of the three models are plotted on top of the transformed signal intensities. Note that the two models “Butterfly (probe type I/sample)” and “Butterfly (probe type II/sample)” together give “Butterfly (probe type/sample)”, which will be used as one method later.

### 3.4. Whole genome sequencing (WGS)

We only used biallic SNPs and alleles that were called with a read depth of at least 25. This resulted in the number of SNPs called shown in Fig. 4.

**Figure 4:**
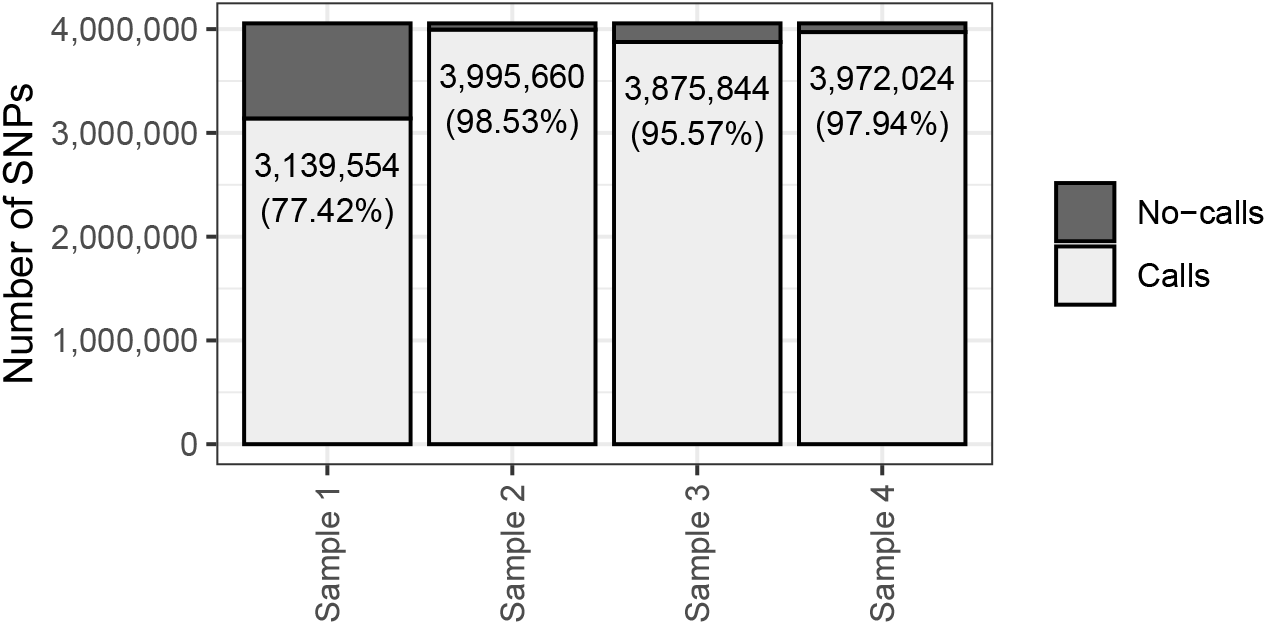
Number of biallelic SNPs called using whole genome sequencing (WGS) with a read depth of at least 25.

Focusing on only SNPs called reliable with WGS, we calculated the concordance rate as we will describe below.

### 3.5. Calling SNPs

The SNP calling is illustrated in Fig. 5 for 400 ng DNA with the “Butterfly (sample)” method and an *a posteriori* probability threshold for NC of 0.8. Fig. 6 shows the homozygous calls for 400 ng DNA without stratifying for individual and show the intensity of the nucleotide in the homozygous call compared to the intensity of the alternative nucleotide (i.e., A for homozygous BB and B for homozygous AA).

**Figure 5:**
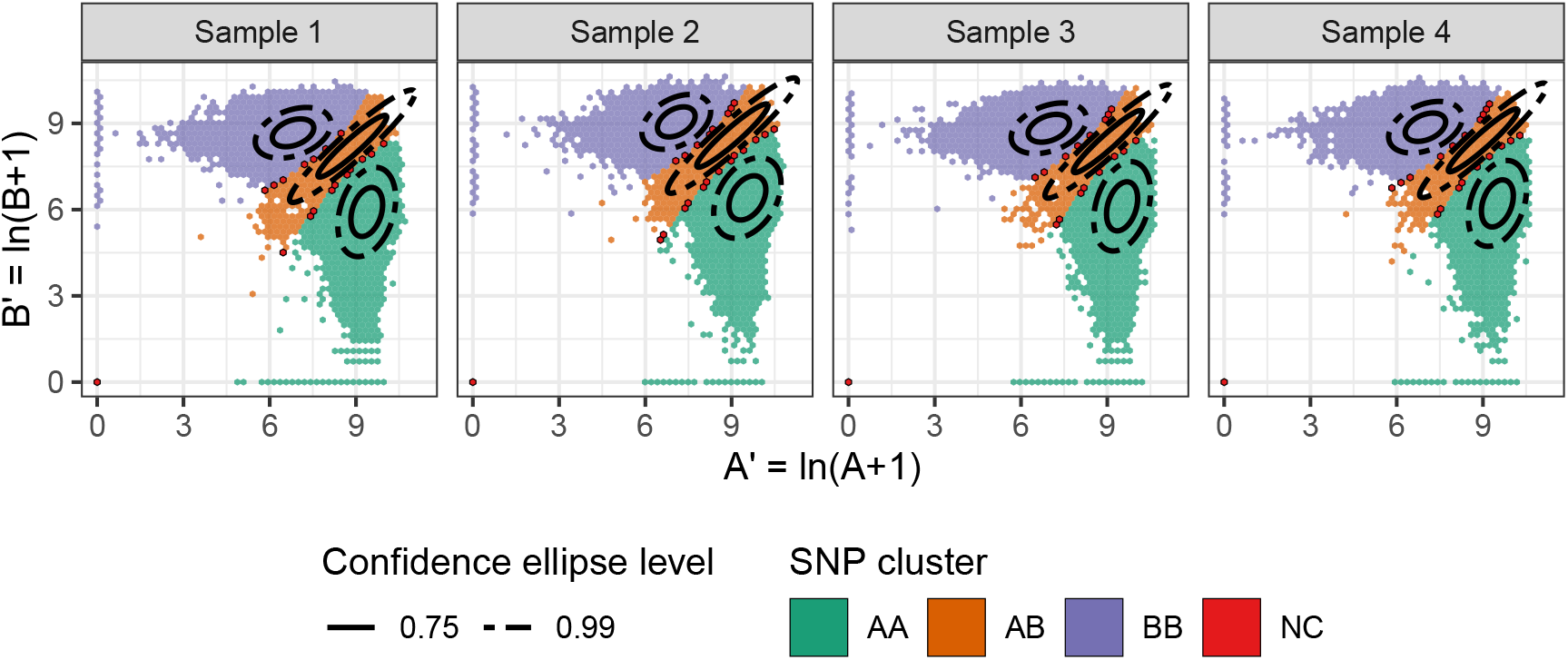
The SNP calling for the “Butterfly (sample)” method illustrated for samples with 400 ng DNA. An *a posteriori* probability threshold of 0.8 was applied for NC. Note that the NCs are in the regions between two SNP groups and at (0, 0).

**Figure 6:**
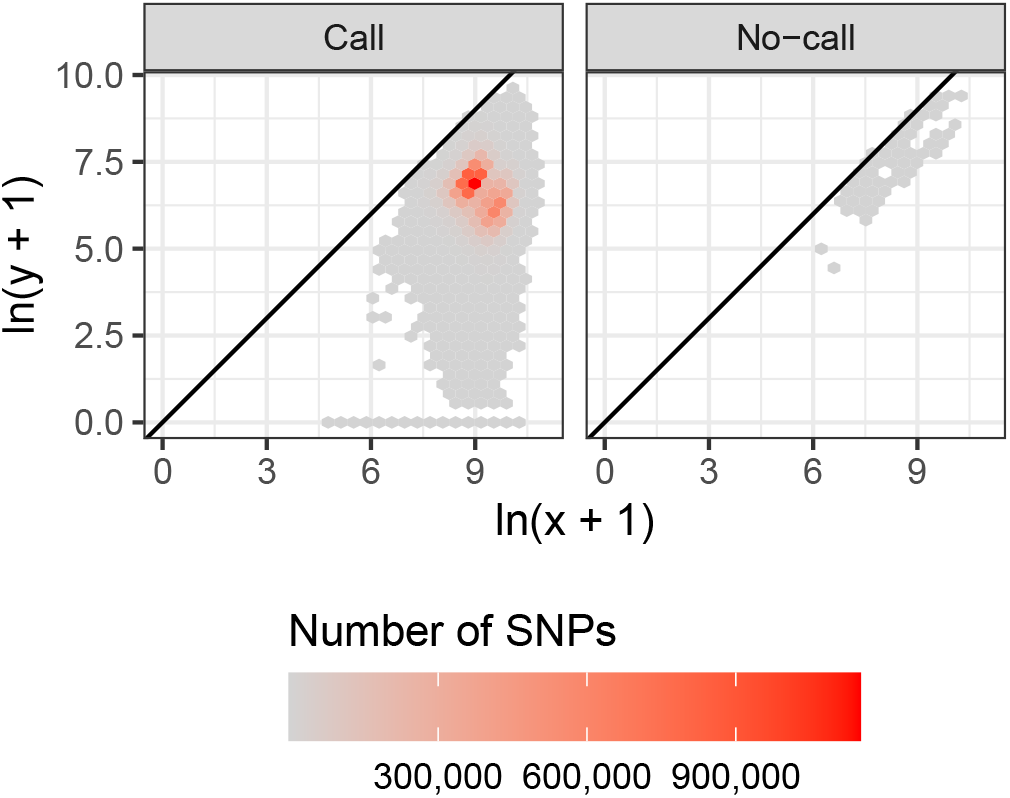
The homozygous SNP calls from Fig. 5 are used (not stratified for individuals) to show the mean intensity, *x*, of the nucleotide in the homozygous call compared to the mean intensity, *y*, of the alternative nucleotide (i.e., A for homozygous BB and B for homozygous AA).

Of the SNPs called by WGS, the number of SNPs called by the other methods are given in Fig. 7.

**Figure 7:**
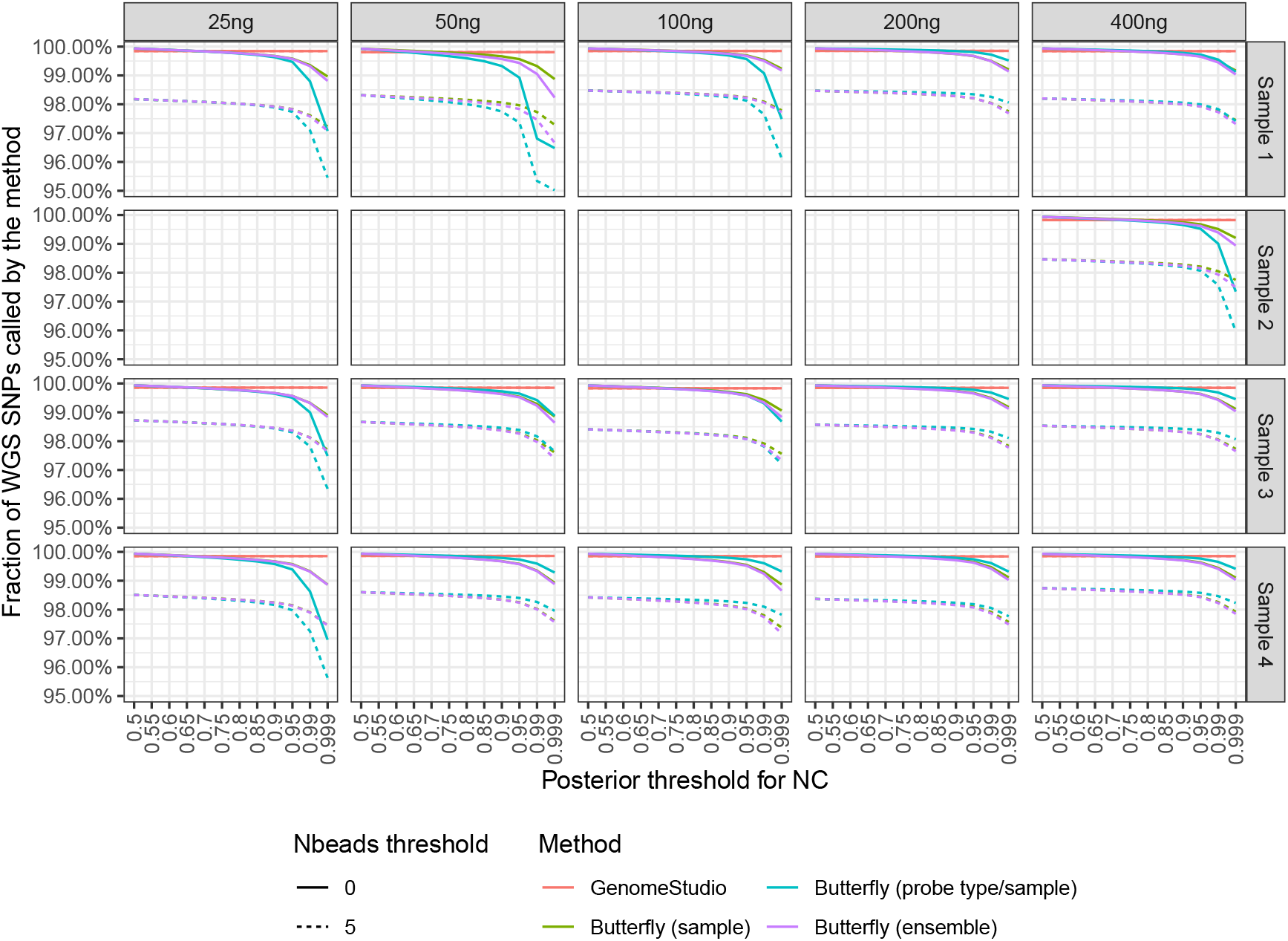
Fraction of SNP calls with the three variations of the butterfly method and GenomeStudio. The butterfly method gave a no-call (NC) if either the maximal *a posteriori* threshold of belonging to a SNP cluster was too low or if the mean signal intensity of either allele A or B (in A/B nomenclature) was based on too few beads (both thresholds shown).

For the SNPs where WGS made a SNP call (i.e., not NC) and when also another method made a call (i.e., not NC), the concordances between the WGS call and the methods are depicted in Fig. 8. This figure shows how reliable a method’s calls are when the NCs are excluded.

**Figure 8:**
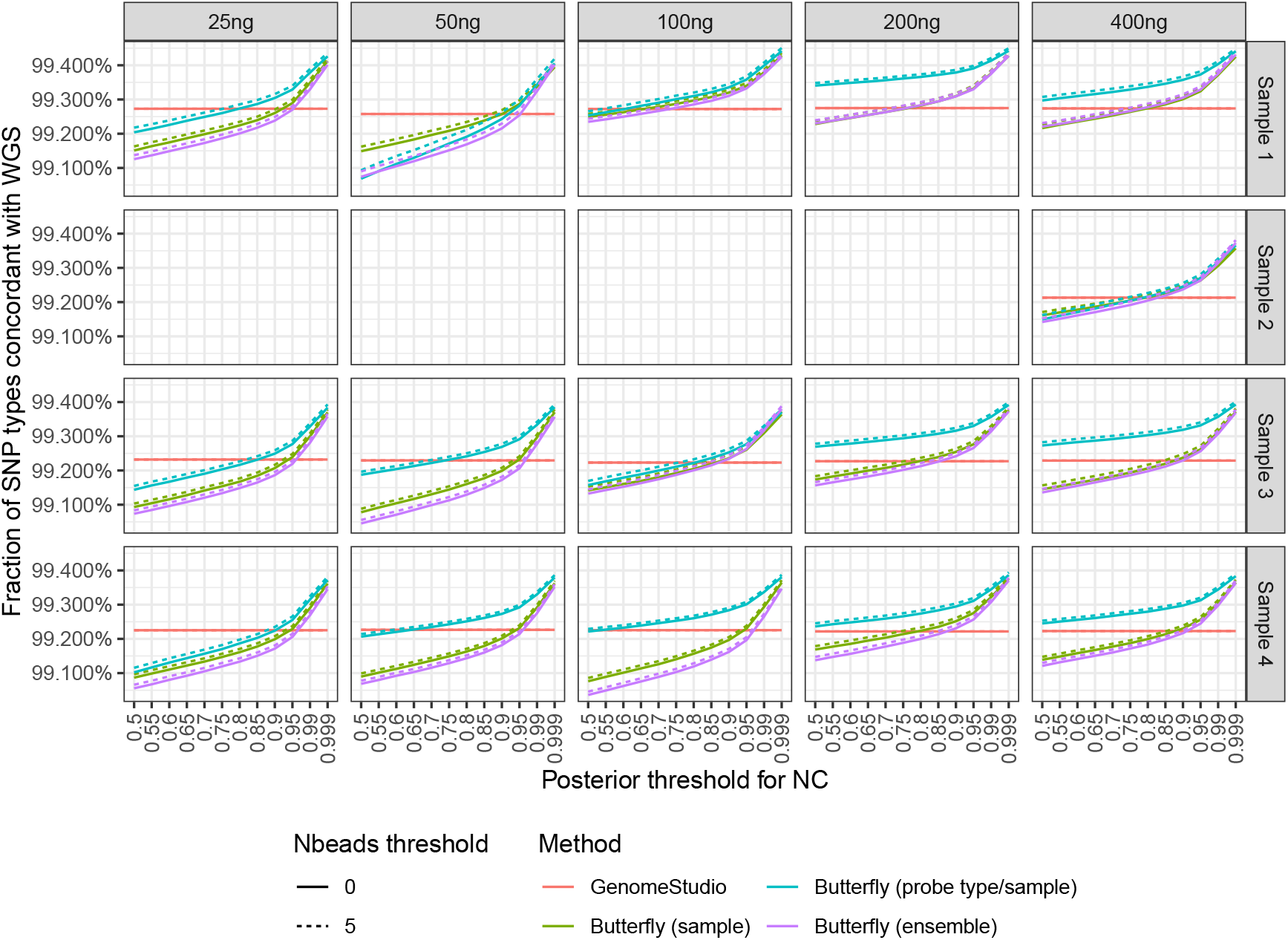
Concordance of SNP calls obtained with WGS to both GenomeStudio and the butterfly methods. When calculating the concordance between WGS and another method, we only focused on SNPs where both methods made SNP calls (i.e., SNPs with NC by any method were excluded).

If one accepts fewer calls by increasing the *a posteriori* probability threshold and the number of beads threshold, the calls will be more reliable (Fig. 7 and Fig. 8).

The importance of the input DNA amount for choosing an *a posteriori* probability threshold can be seen in Fig. 9. For a fixed concordance, the *a posteriori* threshold must be increased the smaller the DNA amount is. The lines did not follow the DNA amount ordering possibly due to saturation and/or quantification error, but the lines of 25 ng and 50 ng DNA were consistently below those of 100 ng to 400 ng DNA.

**Figure 9:**
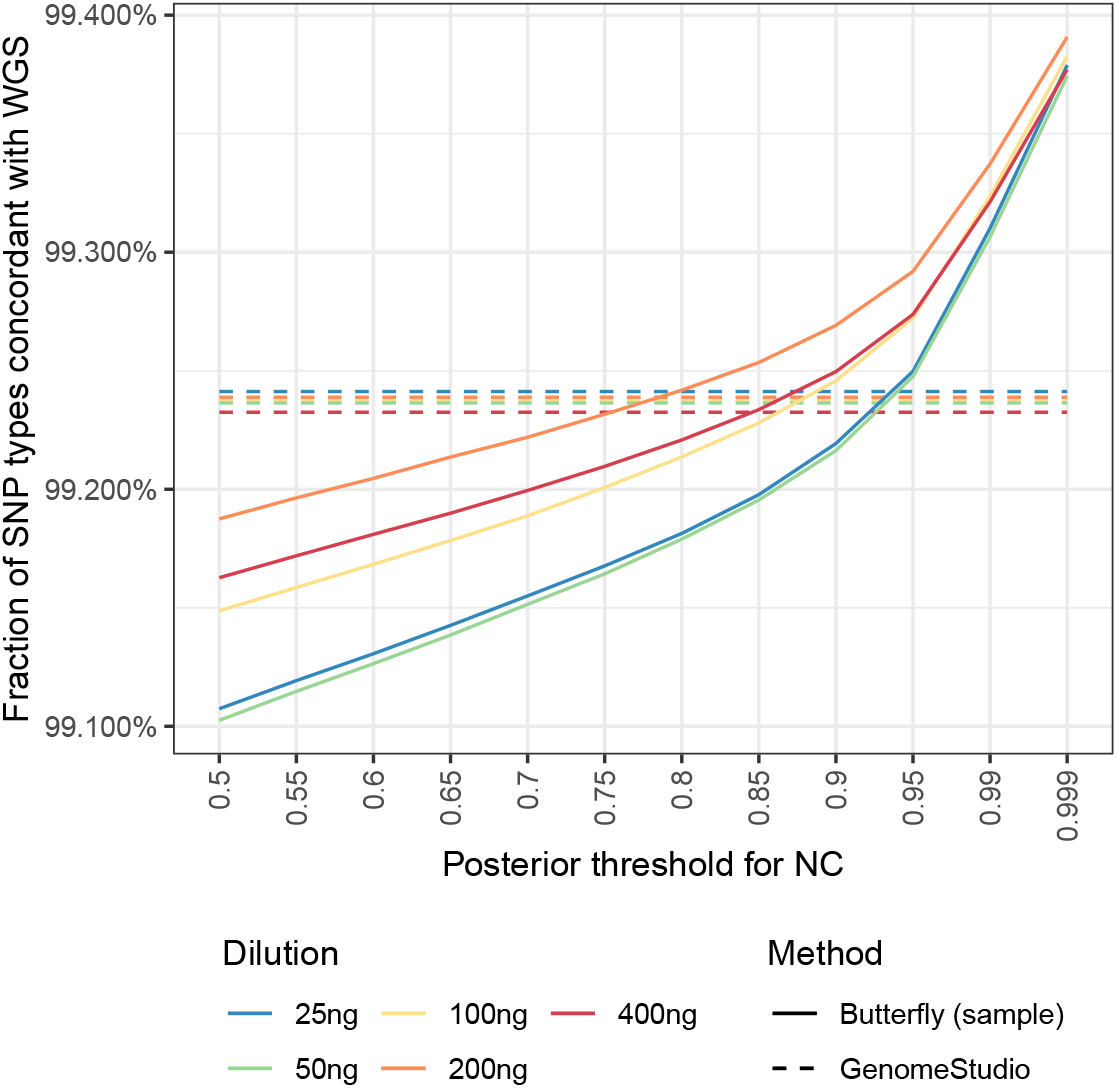
Concordance of SNP calls obtained with WGS to both GenomeStudio and the butterfly (sample) method similar to Fig. 8 (see its caption) aggregated over individuals.

An overview of the discordant calls (excluding no-calls for both WGS and the methods) for 400 ng DNA are shown in Fig. 10.

**Figure 10:**
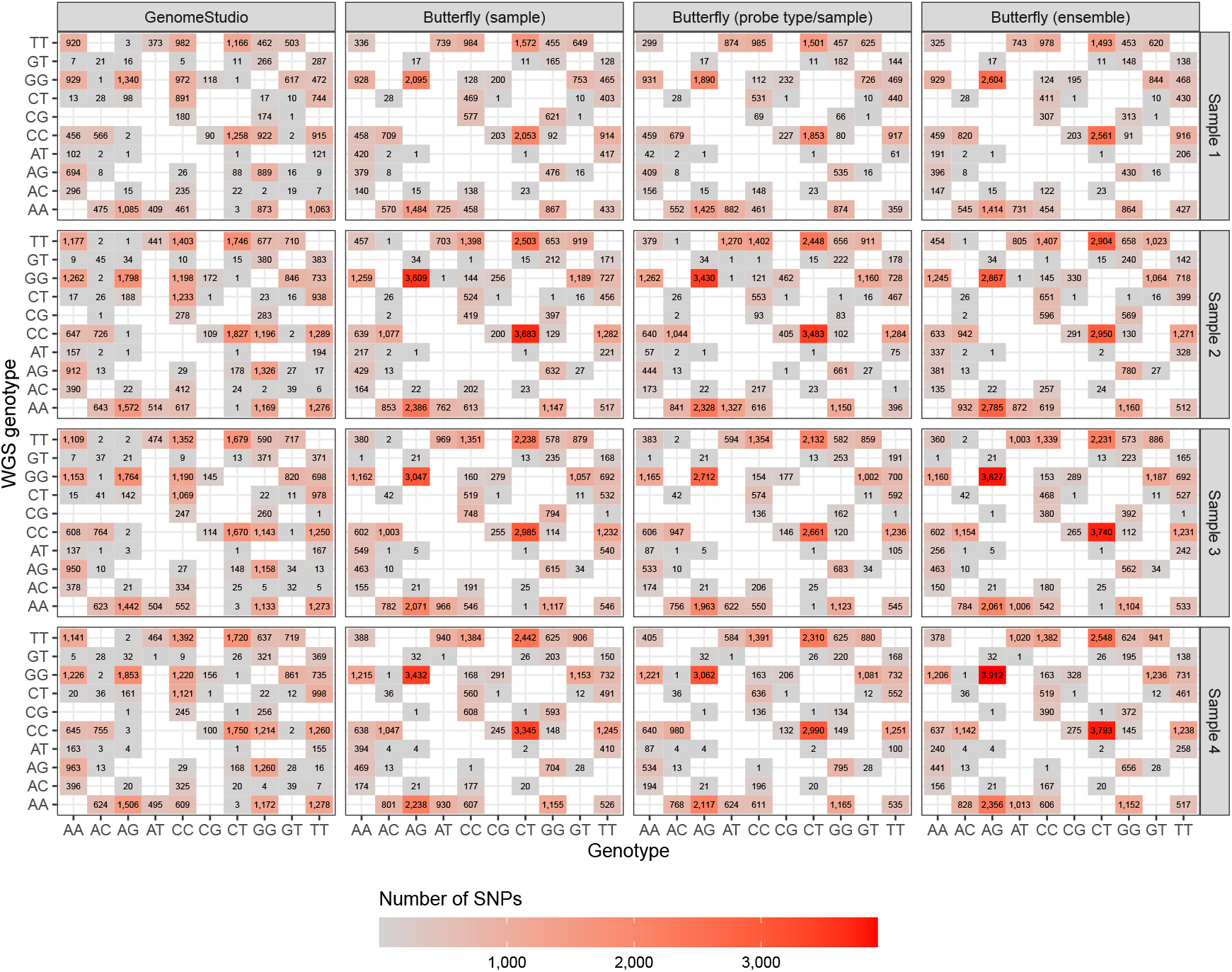
Discordant calls (excluding no-calls) for the samples with 400 ng DNA. The butterfly methods had an *a posteriori* probability threshold of 80% and a number of beads threshold of 0.

Based on Fig. 10, it seems like the butterfly method’s discordancies are due to calling heterozygous when WGS called homozygous and mostly calling AG instead of GG, CT instead of CC, CT instead of TT, and AG instead of AA. A similar pattern is seen with GenomeStudio, but not to the same degree. GenomeStudio made more homozygous discordancies, e.g., AA instead of TT, TT instead of AA, CC instead of GG, etc.

## 4. Discussion

Of the SNPs called with WGS, the butterfly method called more than 99.5% unless high thresholds for *a posteriori* probability and number of beads were used (cf. Fig. 7). For the called SNPs, the concordance between the butterfly method and WGS was 99.0%-99.5% (Fig. 8).

We began with 4,055,428 SNPs (Table 1). Extrapolating from this with the uncertainty involved, we expected the butterfly method to make SNP calls of approximately 4,035,151 SNPs and no-calls for the remaining 20,277 SNPs. Of the called SNPs, 3,994,799-4,010,940 SNPs had reliable calls and 24,211-40,352 SNPs no-calls. This gives a concordant call-rate of all SNPs of around 0.99^2^ = 98%, not taking the uncertainty into account. This emphasises that the numbers are adjustable by the two proposed thresholds (*a posteriori* probability and number of beads), which is easily done using the `R` packages `mclust` [15] and `snpbeadchip` [6].

The importance of the DNA amount and the choice of the *a posteriori* probability threshold can be seen in Fig. 8, which shows that for a fixed concordance, the *a posteriori* threshold must generally be increased for smaller DNA amounts.

Improving the SNP calling is a topic of future research. There are many ways to improve the SNP calling proposed here. Using the WGS calls as reference (with the pitfalls such a decision has), a natural next step of modelling is discriminant analysis based on Gaussianity in a supervised learning setting. Including more explanatory variables also enables more advanced statistical learning methods. In this study, we did not include information about the signal variance, which may improve the SNP calling. Another option is to use probe information like base composition, colour channel, neighbour bases, etc., as explanatory variables/features. This may enable statistical learning methods like multinomial logistic regression, random forests, and deep learning.

